# Transient uncoupling of the Suc-Tre6P-SnRK1 nexus during salt stress associates with biphasic metabolic reprogramming and root plasticity

**DOI:** 10.64898/2026.05.08.723798

**Authors:** Barbieri Giuliano, Parola Rodrigo, Feil Regina, Marianela S. Rodriguez

**Affiliations:** Unidad de Estudios Agropecuarios (UDEA- INTA-CONICET) Camino 60 cuadras km 5.5 X5020ICA, Córdoba, Argentina; Instituto de Fisiología y Recursos Genéticos Vegetales, Ing Agr Victorio Segundo Trippi (IFRGV), Centro de Investigaciones Agropecuarias (CIAP), Instituto Nacional de Tecnología Agropecuaria (INTA), Camino 60 cuadras km 5.5, X5020ICA, Córdoba, Argentina; Max Planck Institute of Molecular Plant Physiology, Am Muehlenberg 1, 14476 Potsdam-Golm, Germany

**Keywords:** Alfalfa, metabolism, root, salt stress, SnRK1, sugar signal, Tre6P

## Abstract

Soil salinization threatens global agriculture reducing yields, yet the metabolic signals controlling salt-sensitive root plasticity in alfalfa remain unclear. We hypothesize that salinity transiently uncouples the sucrose–trehalose-6-P (Tre6P)– Sucrose non-fermenting kinase 1 (SnRK1) nexus, aligning with a biphasic root metabolic response and altered root architecture. Alfalfa seedlings were grown in a hydroponic system and exposed to 200 mM NaCl, with root samples collected from 1 h to 7 d. While primary root growth and biomass remained unchanged, lateral root development was enhanced under salinity. Early response (1 h–1 d) was characterized by reduced carbon metabolites, low Tre6P, increased malondialdehyde, and SnRK1 activation, with a decline in glycolytic and TCA intermediates. During this phase, sucrose was negatively correlated with both Tre6P and SnRK1. Late response (3–7 d) showed a SnRK1 reactivation, Tre6P recovery, and osmoprotectant accumulation, including increased antioxidant capacity (+75% at 3dpt), proline (+178%), and sucrose (+18%) and starch depletion (−57%) at 7dpt respect to control. These metabolic changes coincided with the enhanced lateral root emergence. These findings indicate a two-phase response: early metabolic downscaling with transient Suc–Tre6P–SnRK1 disruption, followed by recovery with Tre6P restoration, SnRK1 reactivation, osmoprotection, and sustained root plasticity under salinity.

**Highlight:** Salinity triggers a temporary metabolic shift in alfalfa roots: plants first conserve energy, then adapt to stress, maintaining lateral root growth and flexible root architecture.

## 1. Introduction

Alfalfa (*Medicago sativa* L.) is a globally important perennial legume valued for its high protein content, yield potential, and the ecosystem services provided by its deep and vigorous root system, including nitrogen fixation, carbon sequestration, and improvements in soil structure and fertility (Wang et al., 2022; Kizildeniz et al., 2024). However, these benefits are increasingly threatened by soil salinization. Although alfalfa is classified as moderately salt-tolerant among legumes (Munns & Tester, 2008), elevated concentrations of sodium (Na□) and chloride (Cl□) severely reduce growth and productivity by imposing osmotic and ionic stress in the rhizosphere (Shelden et al., 2023; Fu et al., 2025).

As the primary interface between plants and saline environments, roots exhibit complex and dynamic responses to osmotic and ionic stress (Duan et al. 2014). Plant responses to salt stress are divided into two temporally distinct phases: Phase I represents the rapid response to increased external osmotic potential, perceived as an initial water deficit, whereas Phase II develops over time as Na□ and Cl□ ions accumulate in plant tissues, leading to ionic toxicity. High salinity typically inhibits overall root development by suppressing meristem cell cycle activity, thereby restricting both primary (PR) and lateral root (LR) growth (West et al., 2004). These architectural alterations, coupled with reduced water and nutrient uptake (Zhao et al., 2023), ultimately cascade into whole-plant dysfunctions, including decreased photosynthetic capacity and inefficient carbon and nitrogen fixation (Farooq et al., 2017; Feng et al., 2025). Nevertheless, plants have evolved stress-induced morphogenic strategies to face these effects. For instance, developmental plasticity under mild stress can trigger LR proliferation, enhancing salt tolerance (Zolla et al., 2010). Similarly, in crops like alfalfa, salinity tolerance varies significantly among cultivars, with resilient genotypes successfully maintaining root growth through superior ion homeostasis. However, the underlying metabolic signaling networks that enable tolerant roots to prioritize and sustain growth under such energy-limiting conditions remain poorly understood.

Carbon availability is a major determinant of root architecture under stress. As heterotrophic sink organs, roots rely entirely on sucrose (Suc) synthesized in aerial tissues and transported via the phloem, as a result, their growth is highly responsive to how the plant allocates its photoassimilates (Lemoine *et al*., 2013). Consequently, research on salinity tolerance has traditionally prioritized shoot responses. However, sustained whole-plant performance under salinity ultimately depends on the ability of roots to adjust their growth and metabolism to cope with systemic carbon limitation. While transient carbon restriction can stimulate PR and LR growth, prolonged deficits severely suppress meristematic activity and overall root development (Belda-Palazón et al., 2020), underscoring the necessity for precise carbon signaling in sink tissues. Central to this regulatory hub is trehalose 6-phosphate (Tre6P), a sucrose-derived signaling metabolite that acts as a highly sensitive indicator of cellular carbon status (Yadav et al., 2014). The Suc–Tre6P regulatory nexus integrates metabolic, hormonal, and developmental signals, translating the availability of photoassimilates into either the promotion or arrest of growth (Figueroa et al., 2016). Specifically, elevated Suc promotes Tre6P accumulation, driving carbon utilization and root growth, whereas Suc depletion reduces Tre6P levels, thereby triggering carbon conservation and repressing growth (Paul et al., 2020). Although the Suc–Tre6P axis is known to play a pivotal role in modulating root meristem activity and LR development (Morales-Herrera et al., 2023), its specific contribution to root architectural adaptation under salinity stress in alfalfa remains an important but unresolved question.

Keeping cellular energy homeostasis under carbon limitation is critically governed by the Sucrose non-fermenting kinase 1 (SnRK1), a highly conserved metabolic sensor that represses energy-consuming anabolic processes and activates catabolic pathways during stress (Baena-González *et al*., 2007). In roots, SnRK1 integrates carbon availability with hormonal and environmental signals to regulate meristem maintenance, LR initiation, and overall developmental plasticity (Crozet *et al*., 2015; Jamsheer *et al*., 2021). Tre6P acts as a primary upstream modulator of this energy-sensing pathway. Under carbon-replete conditions, elevated Tre6P strongly inhibits SnRK1 activity. In contrast, when Tre6P levels fall during salinity-induced carbon restriction, this repression is alleviated, triggering SnRK1 activation to prioritize survival and energy conservation overgrowth. While we recently demonstrated the dynamic and transient behavior of these metabolic signals in salt-stressed alfalfa leaves (Barbieri et al., 2025), how these responses extend to roots, or whether similar transient signaling occurs, remains unclear. Consequently, despite the established central role of this signaling hub, the precise mechanisms by which salinity stress perturbs the coordinated Suc–Tre6P–SnRK1 network to regulate metabolism and architecture in root sink tissues remain largely unresolved.

To address this critical knowledge gap and capture the spatiotemporal dynamics of metabolic reprogramming, we characterized the temporal responses of alfalfa roots to salinity, explicitly differentiating between early response (hours) and late response (days) stress constraints. Focusing on the regulation of the Suc–Tre6P–SnRK1 signaling axis and its impact on root architecture, we integrated LC–MS-based metabolomics with biochemical and molecular profiling. The application of a sudden 200 mM NaCl shock was a deliberate experimental design choice to temporally capture the rapid transition between initial metabolic disruption and subsequent physiological response. This approach maximized the amplitude of the metabolic signals, which would otherwise be difficult to detect under gradual stress imposition. This approach provides, to our knowledge, the first dynamic assessment of SnRK1 activity in alfalfa roots, establishing a comprehensive root–shoot framework for carbon signaling under salinity.

We hypothesize that salinity induces a transient, phase-dependent uncoupling of the canonical Suc–Tre6P–SnRK1 regulatory nexus in root sink tissues. This uncoupling may reflect a differential perception of carbon and energy status during the early stages of stress, enabling a rapid metabolic downscaling that prioritizes survival over the growth. As stress progresses, restoration of Tre6P levels and SnRK1 activity would support a transition toward metabolic acclimation. Under this framework, we propose that temporal modulation of this signaling nexus is associated with the maintenance of root architectural plasticity under salinity.

## 2. Materials and Methods

### Biological Material and experimental setup

Alfalfa seeds of the salt-tolerant Argentinian cultivar Kumen PV INTA (Pecetti *et al*., 2024), a non-dormant with a rest grade of 8 and resistant to *Phytophthora*, anthracnose, and fusarium wilt, were provided by the alfalfa breeding program of the National Institute of Agriculture Technology (INTA).

Seeds were germinated on mesh supports with moist vermiculite placed over 3-liter containers with 5× B&D medium (Broughton and Dilworth, 1971) and continuous aeration. The addition of 0.5 g. L^−1^ NH_4_NO_3_ is a non-nodulation condition to evaluate the effect of salt stress on root development. The Kumen cultivar used in the study was chosen due to its high biomass production, relative salt tolerance, and Na^+^ and Cl^−^ exclusion traits under saline conditions in field trials (Cornacchione and Suarez, 2017) and notable resilience and adaptive responses to salt stress shown in hydroponics (Barbieri *et al*., 2024). Three independent experiments were conducted in a growth chamber under controlled conditions (300 photosynthetically active radiation (PAR) light, 16 h light/8 h dark photoperiod, and 23 ± 2 °C). Salt treatment (200 mM NaCl) was applied 20 days after planting. Root samples were collected at defined time points: 0; 1 and 3 hours post-treatment (hpt); 1; 3- and 7-days post-treatment (dpt). For each treatment, twelve plants were harvested per time point and combined into three biological replicates, each consisting of four roots (Supplementary Fig. S1).

### Ion concentration

Fresh root tissue (100 mg) was ground to a fine powder and dried at 70 °C. The dried material was extracted in 1 ml of 30% (v/v) methanol prepared with Milli-Q water, followed by vortexing for 15 min at room temperature. Samples were centrifuged (6708 g, 10 min), and the supernatant was collected and passed through nylon filters to remove particulates. Ions were determined by HPLC (Shimadzu Prominence Modular HPLC, Kyoto, Japan). Separation was carried out with a Shim-pack IC-C3 column coupled to an IC-C3G guard column, using 3 mM oxalic acid as the mobile phase. Chromatographic runs were performed at 40 °C with a flow rate of 1.2 ml min□¹ for 18 min under non-suppressed conditions, and ions were detected by conductivity. Chromatography was registered and analyzed using LabSolutions (ver. 5.6) software (Pantsar-Kallio and Manninen, 1995).

### Primary metabolite quantification

Alfalfa roots were ground to a fine powder in liquid nitrogen and aliquots (15–20 mg) of frozen powder were extracted with chloroform/methanol as described by Lunn *et al*. (2006). Primary metabolites were measured by LC-MS/MS as described by Lunn *et al*. (2006) with modifications as described by Figueroa *et al*. (2015).

### Soluble Sugars

Root tissue (100 mg) was pulverized in liquid nitrogen and homogenized with 1 ml 80% ethanol (4°C). Following incubation (30°C, 30 min) and centrifugation (12000 g, 10 min), 1 ml acetonitrile was added for purification. After a second spin, 1 ml supernatant was evaporated (90°C, 1 h) and the dried residue dissolved in 700 µL 75% acetonitrile. Sugars were separated via HPLC (Shimadzu) on a ZORBAX amine column using an isocratic mobile phase of acetonitrile:water (81:19) flowing at 1 ml. min□¹. Detection was by refractive index, with quantification against external standards.

### SnRK1 activity assay

SnRK1 activity was measured as previously described (Barbieri et al., 2025). Briefly, 100 mg of fresh root tissue harvested after salt treatment was homogenized in ice-cold extraction buffer supplemented with protease and phosphatase inhibitors. The extracts were clarified by centrifugation (12000 g, 5 min, 4 °C) and desalted using Sephadex G-25 columns. SnRK1 activity was determined by the Universal Kinase Activity Kit (R&D Systems, Minneapolis, MN, United States, EA004) by using AMARA (AMARAASAAALARRR) (Sugden *et al*., 1999) polypeptide as the substrate.

### Immunoblot analysis of SnRK1 T-loop phosphorylation

Phosphorylation of the conserved threonine residue in the activation T-loop of the SnRK1 α-subunit was assessed using a polyclonal antibody recognizing phospho-Thr172 of AMPK (Cell Signaling Technology, Danvers, MA, USA). For immunoblot analysis, 20 μg of soluble extracts were separated by 12% SDS–PAGE and transferred to PVDF membranes. Immunodetection was performed with alkaline phosphatase–conjugated secondary antibodies, and signal development was achieved using an NBT (nitroblue tetrazolium chloride)/BCIP (5-bromo-4-chloro-3’-indolylphosphate p-toluidine salt) substrate system (Invitrogen, #34042).

### RNA isolation, cDNA synthesis, and qRT-PCR analysis

For transcript analysis, root samples (pooled from three plants) were used for gene expression analysis. Total RNA was extracted using TRIzol, and 2 µg of DNase-treated RNA were reverse-transcribed with M-MLV reverse transcriptase (Promega) using oligo(dT) and random primers. qPCR was performed on an iQ5 system (Bio-Rad) with SYBR Green Supermix, following the manufacturer’s instructions. Expression levels were analyzed using the 2^DΔΔCt^ method (relative to housekeeping and control) according to Livak and Schmittgen (2001). A gene fragment encoding alfalfa ubiquitin (UBQ, AW686873) was used as an internal control to normalize the amount of template cDNA, consistent with the reference gene used by Kakar et al. (2008). The gene-specific primer pairs employed for the detection of alfalfa transcripts of *SnRK1.1* (MS.gene86051.t1), SENESCENCE-ASSOCIATED PROTEIN 5 (*SEN5*, Medtr2g079670.3), and ASPARAGINE SYNTHETASE1 (*ASN1*, Medtr3g464580.1) are provided in Supplementary Table S1. The specific primers were designed by homology analysis between expressed alfalfa sequence tags and public databases. Expression analysis for each gene was repeated three times to ensure technical reliability.

### Statistical Analysis

All experiments were performed with at least three biological replicates. Statistical significance among means was assessed by analysis of variance (ANOVA), followed by Tukey’s post hoc test (*P* < 0.05). For pairwise comparisons, a parametric Student’s t-test was used. The complete metabolite dataset was used to generate a principal component analysis (PCA) biplot. Data processing and statistical analyses were conducted using InfoStat software (Di Rienzo et al., 2019). Pearson correlation analyses were performed using GraphPad Prism version 10.0.0 for macOS (GraphPad Software, Boston, MA, USA).

## 3. Results

### Lateral root increase in salt-tolerant alfalfa cultivar under saline conditions

Root system architecture is highly plastic and undergoes dynamic remodeling under abiotic stress through changes in key traits such as root length, branching, and biomass (Zou et al., 2021). In alfalfa, salt treatment led to a significant increase in LR development since 3 dpt (Fig. 1B). In contrast, PR length (Fig. 1C) and root dry biomass (Fig. 1D) remained unchanged compared to control plants.

**Fig. 1.**
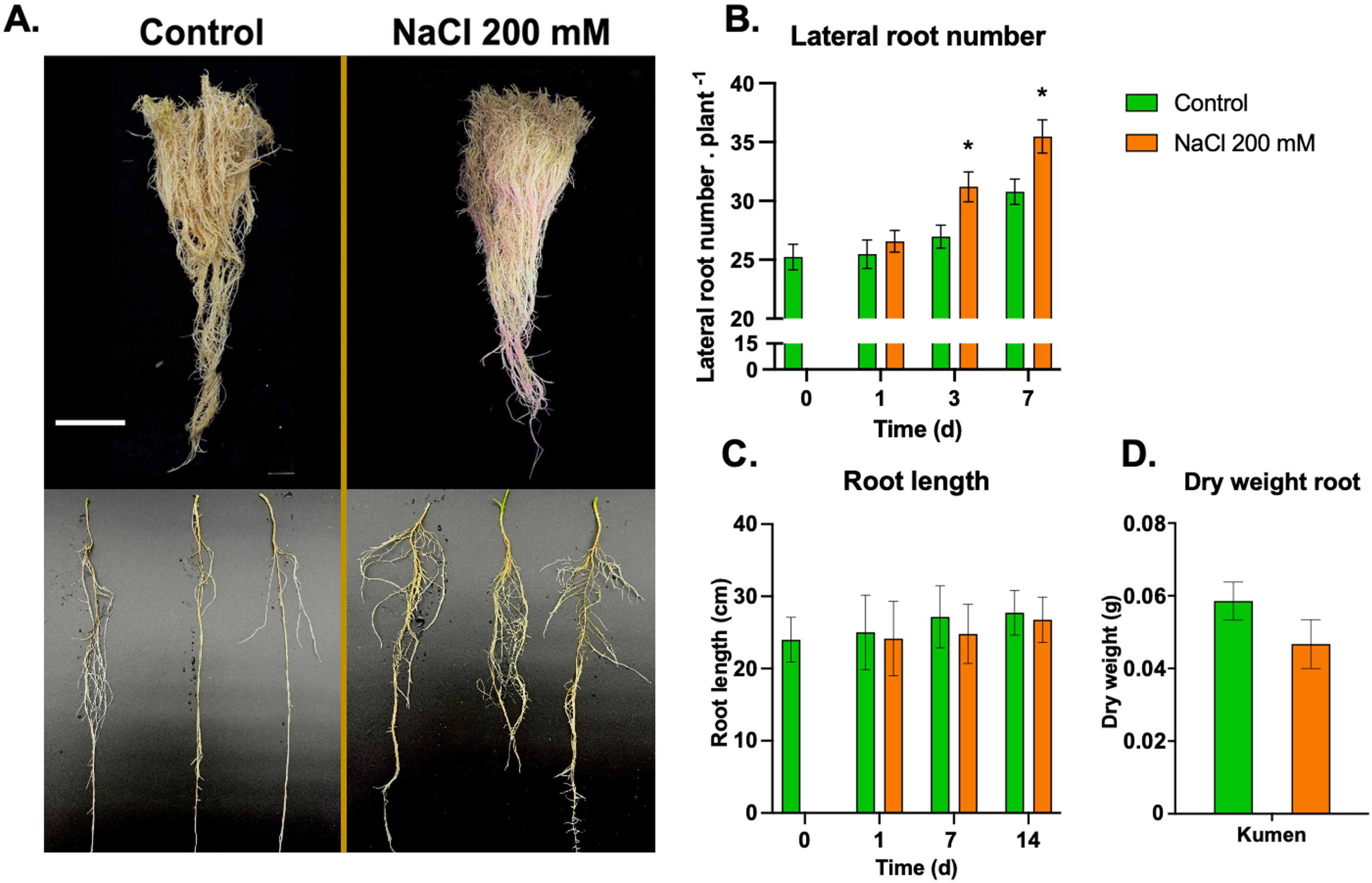
Effects of salt stress on alfalfa root. A. Representative images from three independent experiments showing control (left) and 200 mM NaCl-treated (right) alfalfa root systems (n = 60). B. Lateral root number. C. Root length measured at 0, 1, 7, and 14 days post-treatment. D. Root dry weight after 14 days under control and 200 mM NaCl conditions. Green bars represent control plants and red bars represent NaCl-treated plants. Asterisks indicate significant differences between salt-treated plants and their respective controls, based on three independent experiments (n = 105), analyzed by ANOVA followed by Tukey’s post hoc test (*P* < 0.05).

Ion profiling revealed a rapid and sustained accumulation of Na□ in roots, detectable as early as 1 h post-treatment (hpt) and persisting throughout the experimental period (Supplementary Fig. S2A). The Na□:K□ ratio exhibited a biphasic pattern, characterized by an initial increase at 1 hpt, followed by a gradual decline of 74% by 3 dpt, and a subsequent rise at 7 dpt (Supplementary Fig. S2C).

### Metabolic shift at 1 day post-treatment in response to salt stress in alfalfa roots

Since LR exhibited a significant increase under salt stress in the salt-tolerant alfalfa cultivar evaluated, we investigated the time-dependent metabolic changes occurring in roots during stress exposure. Metabolomic analyses offer a powerful approach to uncovering temporal dynamics of stress responses, providing detailed insights into the modulation of metabolic intermediates involved in glycolysis, the TCA cycle, and polysaccharide biosynthesis (Li et al., 2023).

We observed a significant accumulation of succinate and a decrease in pyruvate at 1 hpt, while at 3 hpt all the metabolites were stabilized and just Tre6P dropped until 1 dpt. By 1 dpt, marked reductions were observed in most glycolytic intermediates, including glucose-6-phosphate (Glc6P), glucose-1,6-bisphosphate (Glc1,6BP), fructose-6-phosphate (Fru6P), fructose-1,6-bisphosphate (FBP), phosphoenolpyruvate (PEP) and 3-phosphoglycerate (3PGA); and in tricarboxylic acid (TCA) cycle intermediates such as citrate, cis-aconitate, isocitrate, fumarate and malate. Concomitantly, proline accumulated in salt-treated roots and remained elevated throughout the experimental period. At 3 dpt, Glc1P and galactose-1-phosphate (Gal1P), metabolites involved in polysaccharide biosynthesis, were significantly reduced. By 7 dpt, UDP-glucose (UDP-Glc) and UDP-Gal, key precursors for cell wall biosynthesis, also showed a significant decrease, accompanied by a reduction in Glc6P and starch content (Fig. 2).

**Fig. 2.**
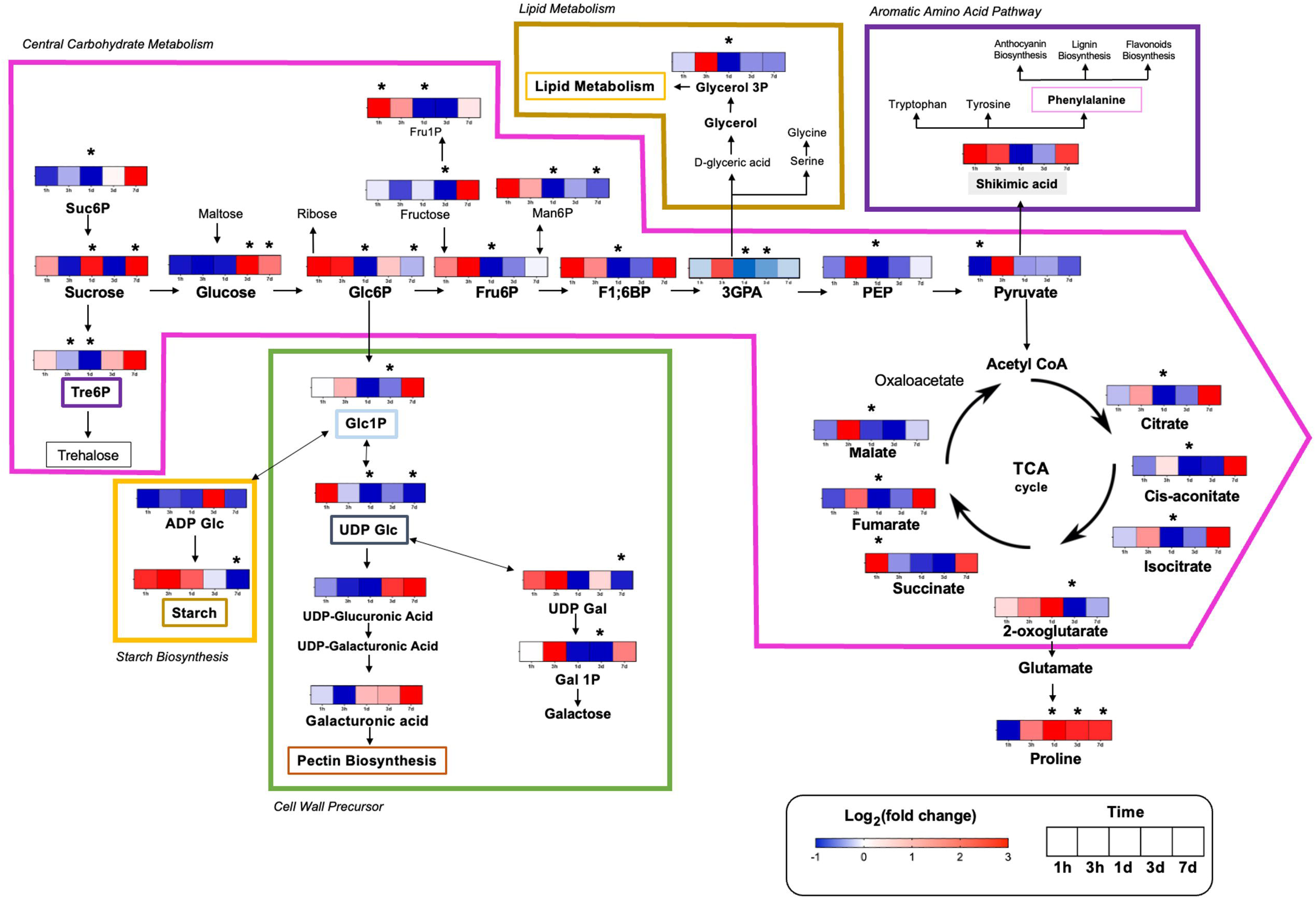
Schematic diagram of response metabolites of alfalfa roots under salt stress. Heat map colors indicate the log2-transformed fold change in metabolite levels in stressed plants relative to controls. Blue denotes reduced metabolite levels, white indicates no variation, and red reflects increased levels. Each cell corresponds to a specific sampling time (1 h, 3 h, 1 d, 3 d, and 7 d). Asterisks mark statistically significant differences between salt-treated plants and their corresponding controls. These results are based on four independent experiments, each comprising four plants per treatment combined into a single sample (n = 16), and were evaluated using ANOVA followed by Student’s t-test (*P* < 0.05).

To identify the metabolites contributing most strongly to the temporal separation among samples, we performed a Variable Importance in Projection (VIP) analysis based on PLS-DA models (Fig. 3). Under salt stress, proline and Glc displayed the highest VIP scores (VIP > 1.5), indicating that these metabolites were among the most influential variables driving the discrimination between time points (Fig. 3A). In control roots, starch and proline showed the highest VIP values (VIP > 1.5), reflecting their contribution to temporal variability under non-stress conditions (Fig. 3B).

**Fig. 3.**
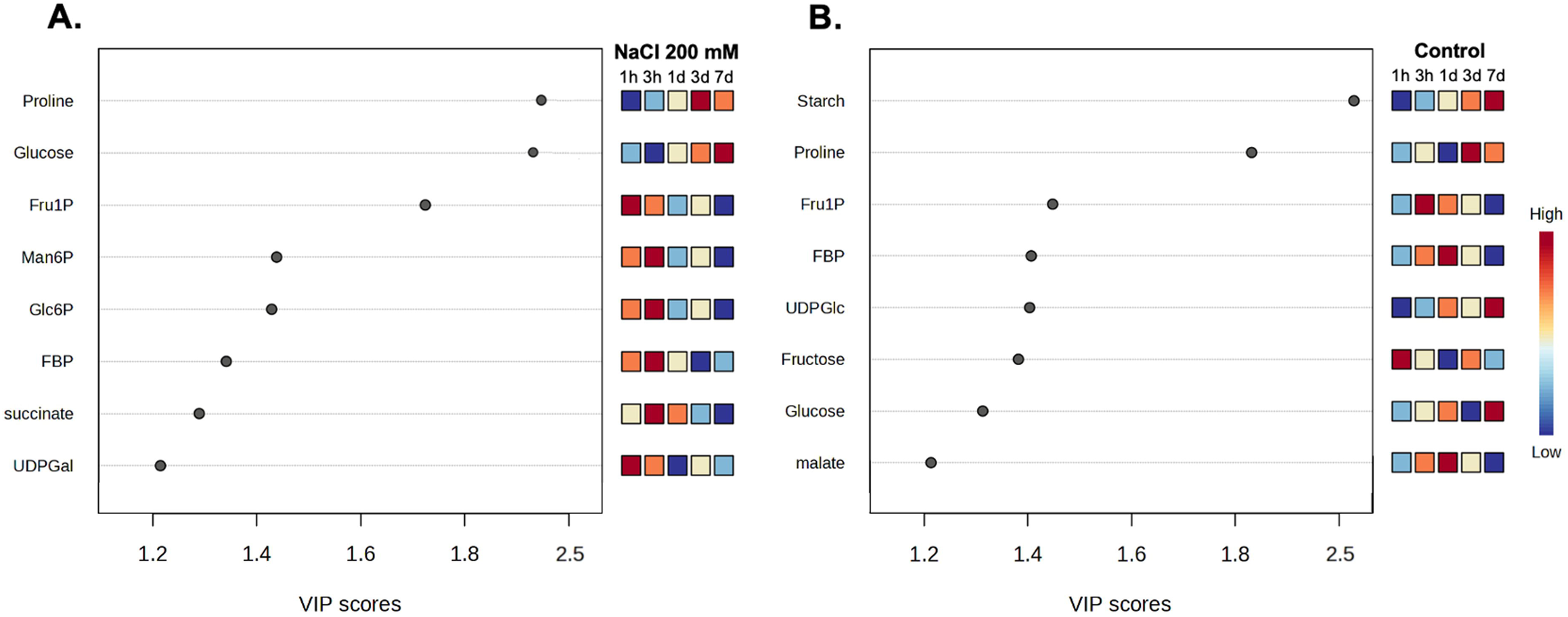
Temporal metabolic profiles of alfalfa roots under salinity and control conditions. A. Roots treated with 200 mM NaCl. B. control roots. Variable Importance in Projection (VIP) scores from Partial Least Squares Discriminant Analysis (PLS-DA) models based on treatment (Control vs NaCl) identify the eight most influential metabolites (VIP > 1.2) contributing to discrimination among time points (1 h, 3 h, 1 d, 3 d, 7 d). The heatmap shows relative metabolite abundances, with red indicating higher levels, blue lower levels, and pale yellow intermediate levels relative to the mean across all time points.

Integration of VIP scores with the associated heatmap patterns revealed a clear biphasic metabolic response to salinity. An early response (1 hpt to 1 dpt) was characterized by transient accumulation of specific intermediates such as Fru1P and Glc, followed by a broad decline in glycolytic and TCA cycle metabolites, including FBP, PEP, 3PGA, and organic acids. This pattern suggests a rapid metabolic adjustment consistent with a shift toward reduced carbon flux and energy conservation. In contrast, a later response (3–7 dpt) was marked by the progressive accumulation of proline and the sustained depletion of nucleotide sugars such as UDP-Glc and UDP-Gal, indicating a reallocation of carbon toward osmoprotective functions at the expense of biosynthetic processes. Together, these results indicate that the metabolites identified by VIP analysis not only contribute to temporal discrimination but also display coordinated changes consistent with a shift in carbon allocation and metabolic activity during stress progression.

### Transient uncoupling of the Suc-Tre6P-SnRK1 nexus in alfalfa roots under salinity

To further elucidate how metabolic signals are integrated under these conditions, we examined the sucrose–Tre6P–SnRK1 regulatory nexus, a central module controlling carbon allocation and root development. SnRK1 represents a central component of this regulatory framework, functioning as a key integrator within the carbon and energy signaling network (Peixoto and Baena-González, 2022). SnRK1 activity was assessed using three complementary approaches to capture both biochemical activity and *in vivo* regulatory outputs. First, *in vitro* kinase activity was measured using a universal kinase assay (Fig. 4C). Second, the phosphorylation status of the conserved T-loop domain was evaluated as an indicator of SnRK1 activation state (Polge and Thomas, 2007) (Supplementary Fig. S3A). Third, transcript levels of the SnRK1-responsive marker gene *MsSEN5* (*Medtr2g079670.3*), homologous to *SEN5* (*AT3g15450*) in Arabidopsis, and *MsASN1* (*Medtr3g464580.1*), homologous to *DIN6* (*AT3G47340*) in Arabidopsis, were analyzed as a proxy for SnRK1 activity *in vivo*; these genes has been widely used as a sensitive readout of SnRK1-dependent transcriptional responses (Baena-González et al., 2007; Ramon et al., 2019) (Supplementary Fig. S3B).

**Fig. 4.**
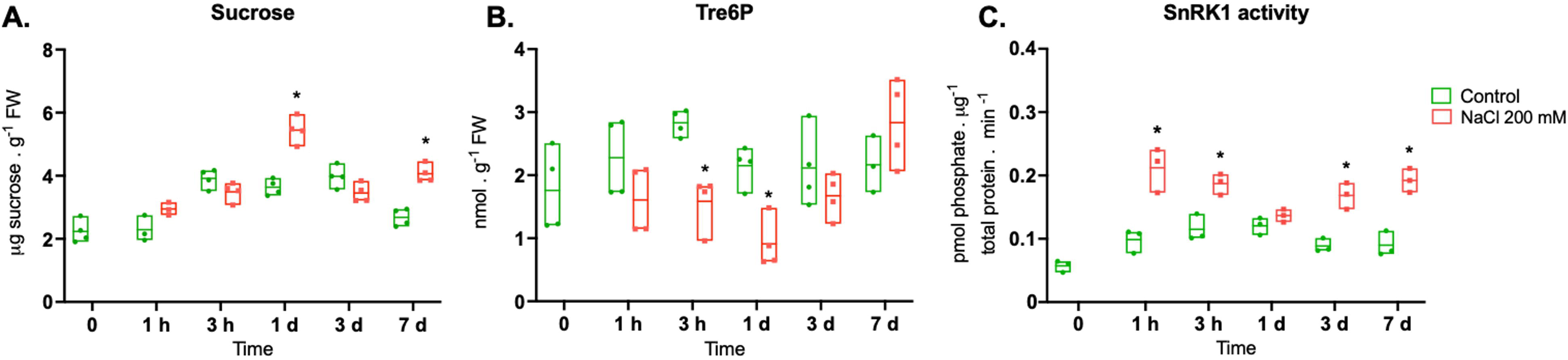
Suc-Tre6P-SnRK1 in alfalfa roots during NaCl stress. Graphs show results for roots from control (green) and treated with NaCl 200 mM (red) grown for 20 days in 0.5X B&D medium. A. Suc concentration measured by HPLC. B. Trehalose-6-phosphate (Tre6P) concentration measured by LC-MS/MS. C. SnRK1 activity measured with universal kinase kit. Asterisks indicate significant differences between plants under saline treatment and their respective controls, based on four independent experiments with four plants per treatment, grouped into one pool (n=16), analyzed using ANOVA followed by Tukey’s test (p < 0.05).

Under salt stress, Suc levels increased significantly at 1 and 7 dpt, +49.7% and +51.4%, respectively, respect to control (Fig. 4A). From a kinetic point of view, Suc level increased by +85,3% between 1 hpt to 1 dpt, whereas Tre6P content exhibited a pronounced decline by −43.7% decrease between 1 hpt to 1 dpt (Fig. 4B). Given that Tre6P has been reported to inhibit SnRK1 activity in sink tissues (Zhai et al., 2018), this decrease is consistent with the overall increase in SnRK1 activity observed across most time points, except for 1 dpt (Fig. 4C). Consistent with these measurements, both T-loop phosphorylation and the expression of the SnRK1-responsive marker genes (MsASN1 and MsSEN5) showed temporal profiles that largely paralleled the kinase activity assay during the early response of the stress response (Fig. 4C; Supplementary Fig. S3B). Interestingly, MsASN1 and MsSEN5 exhibited distinct expression dynamics, with MsASN1 declining earlier than MsSEN5. This divergence highlights that transcriptional markers may differ in their temporal sensitivity and reinforces the need for caution when using individual gene expression readouts as proxies for SnRK1 activity. Together, this data indicate that there are two different phases, that could be related to an early response and the other related to acclimatation response.

The correlation analysis exhibits differences between the Suc-Tre6P-SnRK1 interaction dependent on the phase with a strong inverse relationship between Suc and SnRK1 (r = −0.88, *P*<0.0002), as well as between Suc and Tre6P (r = −0.63, *P*<0.02). These results are consistent with a temporary uncoupling of the canonical Suc–Tre6P–SnRK1 module during the early stress phase (Fig. 5). In this context, we operationally define uncoupling as a temporal mismatch between sucrose levels, Tre6P abundance, and SnRK1 activity, deviating from their canonical regulatory relationships described under non-stress conditions.

**Fig. 5.**
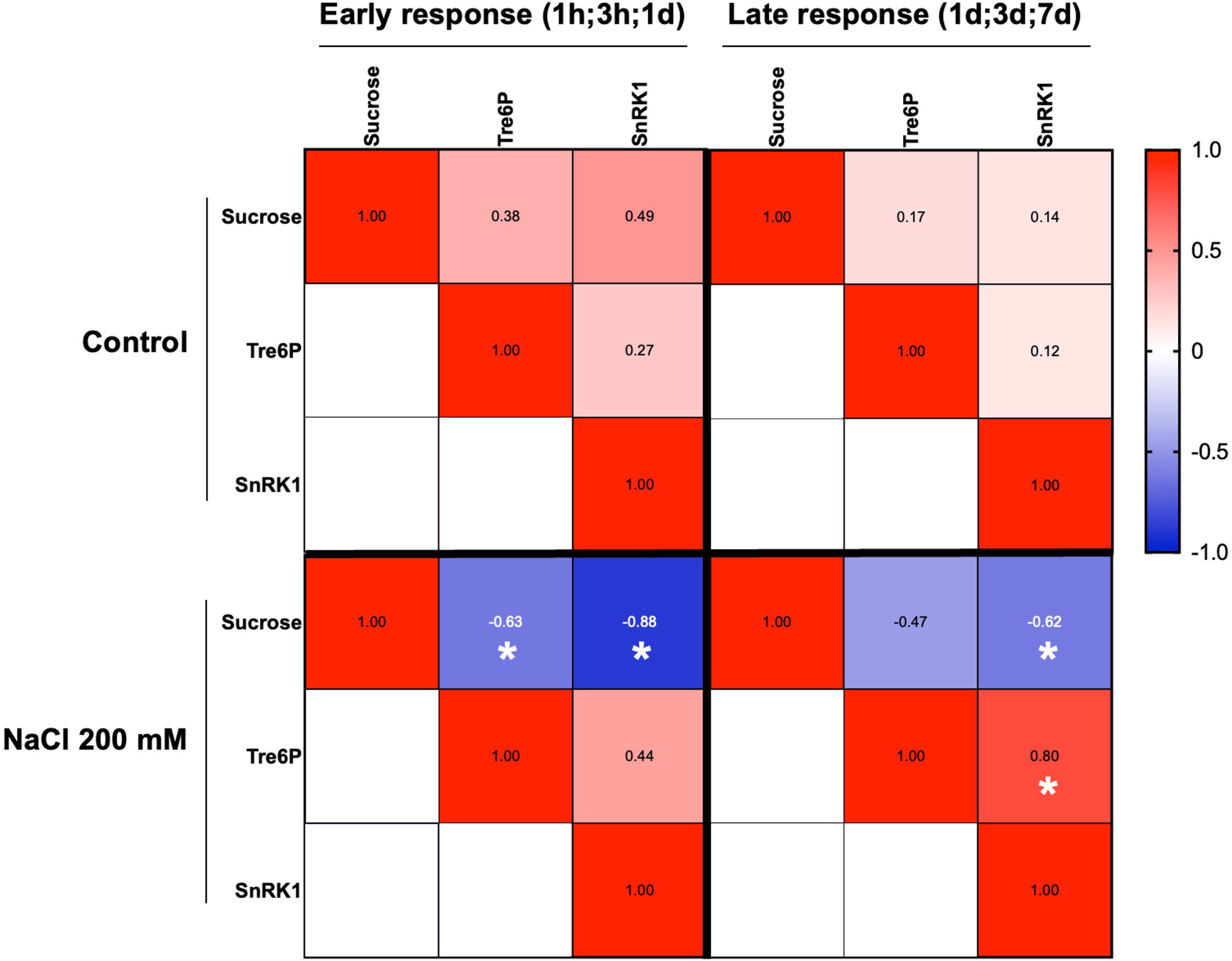
Correlation analysis of Suc, Tre6P, and SnRK1 under control and salt stress conditions. Heatmaps display the Pearson correlation coefficients for metabolite and enzyme activity levels across early response (1h, 3h, 1d) and late response (1d, 3d, 7d) time points. Separate matrices are shown for control and salt-stressed plants. Red cells indicate positive correlations, blue cells indicate negative correlations, and color intensity corresponds to the magnitude of the coefficient. Asterisks indicate a statistically significant correlation (p < 0.05).

### Oxidative status and metabolic changes during late response

To further characterize the responses occurring during salt stress, we examined parameters associated with oxidative status and metabolic adjustments (Fig. 6). In roots, salinity induced a marked modulation of redox-related traits. Total antioxidant capacity, estimated using the FRAP (Ferric Reducing Antioxidant Power) assay as a proxy for global reducing potential, increased significantly at 3 dpt in plants exposed to 200 mM NaCl compared with their respective controls. In contrast, lipid peroxidation, assessed by malondialdehyde (MDA) content, was elevated at early points, with significantly higher levels detected at 1 and 3 hpt under saline conditions. These results indicate a temporal shift from an initial oxidative perturbation to a more balanced redox state during the late response.

**Fig. 6.**
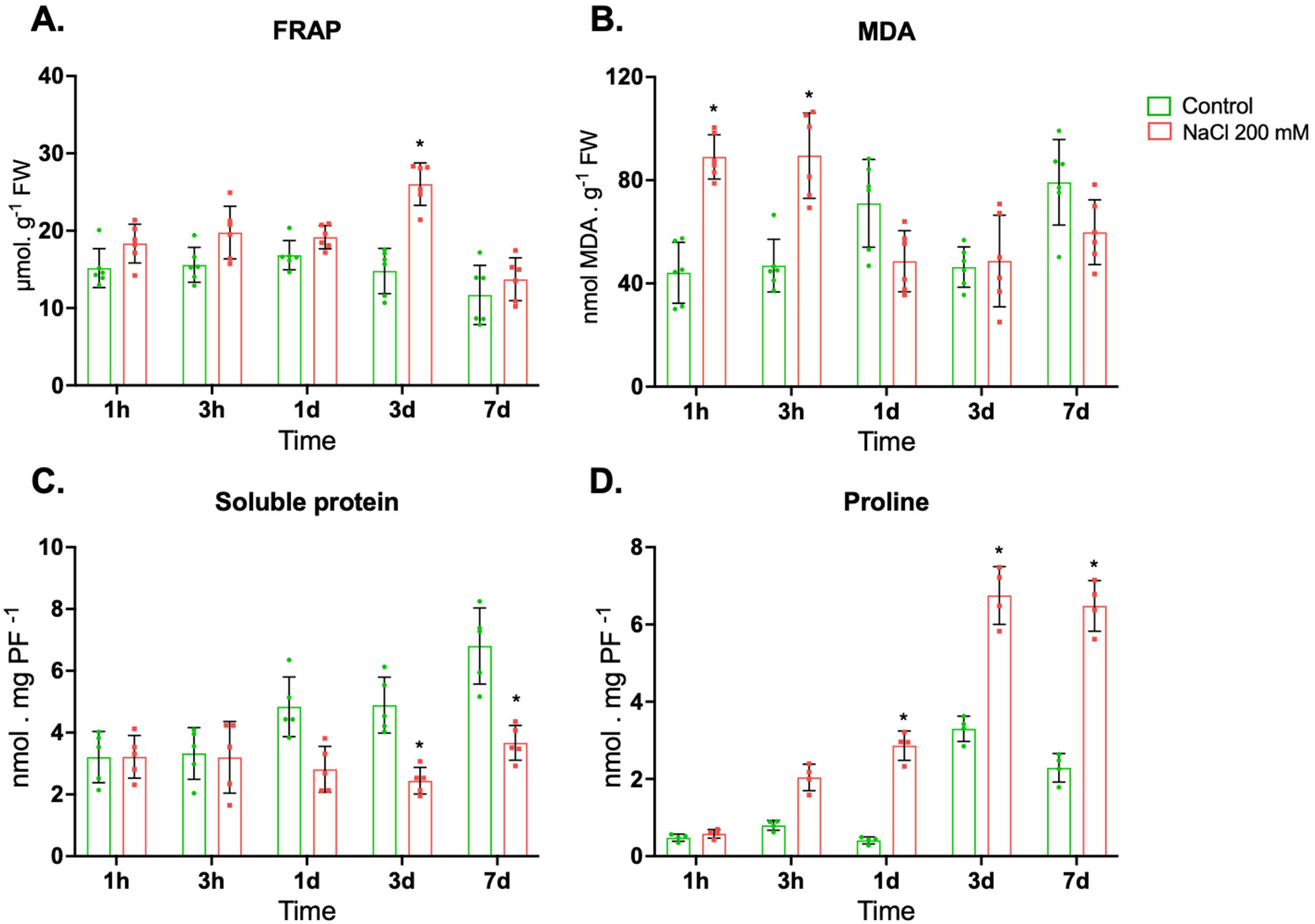
Biochemical dynamic characterization in alfalfa roots under salt stress. A. Ferric reducing antioxidant power (FRAP), B. malondialdehyde (MDA), C. Soluble protein, and D. Proline contents were measured in roots of control plants (green) and plants subjected to 200 mM NaCl (red). Bars represent mean values ± SD. Individual data points are shown. Asterisks indicate significant differences between saline-treated plants and their respective controls at each time point, based on six independent experiments with four plants per treatment pooled per experiment (n = 24), analyzed by ANOVA followed by Tukey’s post hoc test (p < 0.05).

This transition was accompanied by a reconfiguration of carbon and nitrogen metabolism, characterized by reduced levels of UDP-Glc, consistent with alterations in cell wall metabolism. In parallel, total soluble protein content showed a significant decrease at 3 and 7 dpt in salt-treated roots, suggesting enhanced protein degradation under prolonged stress. Conversely, proline content increased significantly from 1 dpt onwards and remained elevated until the end of the experiment, consistent with its role as an osmoprotectant and stress-responsive metabolite, supporting the establishment of a metabolically adjusted state associated with sustained stress tolerance.

### Temporal dynamics of the root metabolome and SnRK1 activity under salt stress

To gain a comprehensive understanding of metabolic reprogramming and evaluate the temporal dynamics in alfalfa roots under salinity, a principal component analysis (PCA) was performed independently for salt-treated (200 mM NaCl) and control plants (Fig. 7A, B). The first two principal components explained 70.2% (PC1 42.4%, PC2 27.8%) and 77.3% (PC1 50.8%, PC2 26.5%) of the total variance in salt-stressed and control roots, respectively. The biplots revealed divergent temporal trajectories for SnRK1 activity and the root metabolome. In salt-stressed roots, SnRK1 activity was linked to the early response (1 hpt) but inversely correlated with the metabolic shift observed at 1 dpt (Fig. 7A). This time point was also characterized by a distinct accumulation of TCA cycle intermediates, including isocitrate, fumarate, and malate. Conversely, our data supports a model where later responses to stress (7 dpt) were primarily associated with metabolites related to osmoprotection (proline) and cell-wall composition (galacturonic acid).

**Fig. 7.**
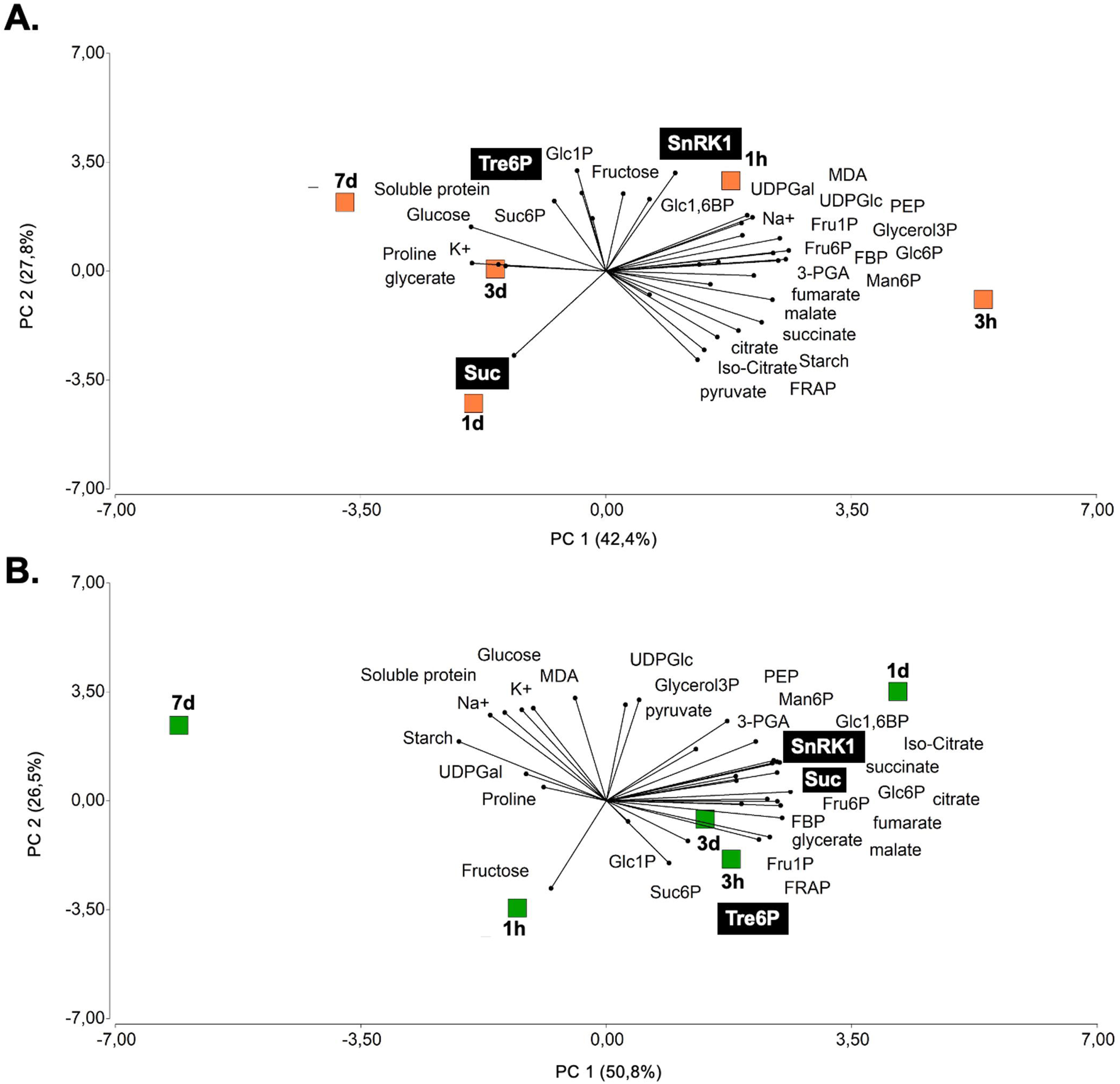
Principal Component Analysis (PCA) biplot obtained from the first and second principal components A. Roots of NaCl 200 mM treated plants; B. control roots. Results are from four independent experiments with a pool of three roots per experiment.

Under control conditions, the metabolic trajectory exhibited a markedly different pattern (Fig. 7B). Notably, the vectors for sucrose and SnRK1 activity were closely aligned, indicating a positive association during the evaluated time course, whereas Tre6P did not strongly associate with the main temporal clusters. Furthermore, TCA-related metabolites in control roots co-segregated with SnRK1 activity and were most closely associated with the metabolic shift around 1 dpt rather than with early developmental time points such as 3 hpt.

## 4. Discussion

Roots serve as the primary interface for sensing and adapting to abiotic soil stresses, making their morphological plasticity a central determinant of plant survival (Rao et al., 2021). In alfalfa, salt tolerance is closely linked to root architectural responses; for instance, highly tolerant cultivars maintain robust LR emergence under saline conditions (Yu et al., 2021). Because these architectural traits are functionally correlated with water and nutrient uptake capacity (Wang et al., 2007; Arif et al., 2019), our results confirm that the salt-tolerant alfalfa cv. Kumen employs a similar adaptive strategy. While overall root biomass and PR elongation remained remarkably stable under salinity, we observed significant stress-induced LR development (Fig. 1). This adaptive response preserves root architectural plasticity and contrasts sharply with the behavior reported in other species, such as rice, which exhibits severe LR reductions even in salt-tolerant cultivars (Ijaz *et al*., 2019). Ultimately, the stability of the alfalfa root system aligns with the broader physiological consensus that root development in this species is significantly less inhibited by salinity than shoot growth, likely ensuring systemic stress acclimation (Cornacchione and Suárez, 2017; Bertrand et al., 2020; Barbieri et al., 2025).

Maintaining root morphological plasticity under stress requires rapid metabolic reprogramming. To capture the signaling events associated with the time-course acclimation, the application of a sudden 200 mM NaCl treatment in this study was specifically designed to expose the rapid, transient metabolic changes that occur immediately after stress perception. This “salt shock” enhances temporal resolution and synchronizes cellular responses, providing a mechanistic, rather than quantitative, framework relative to field conditions, where gradual salinity would likely elicit attenuated dynamics. Even so, defining this response establishes a baseline for how roots overcome sink limitation to prioritize resource allocation and survival. Together, the data support a systemic acclimation strategy in which metabolic reprogramming aligns with developmental plasticity, hormonal control, and resource allocation to sustain root function. The temporal coupling of metabolic shifts with increased lateral root development points to a functional link between energy signaling and plasticity, though direct causality remains to be demonstrated.

The root acclimation is intrinsically linked to whole-plant source–sink dynamics. As we recently demonstrated, salt stress induces significant chloroplast alterations and progressive Suc accumulation in alfalfa source leaves up to 1 dpt (Barbieri *et al*., 2025). Concurrently, ion profiling revealed a rapid and sustained Na□ accumulation alongside biphasic Na□:K□ dynamics, indicating an early ionic imbalance, a transient recovery of ion homeostasis, and a subsequent shift under prolonged stress.

The transient carbon retention in mature leaves observed by Barbieri *et al*. (2025), likely restricts photoassimilate transport to heterotrophic roots, triggering a biphasic metabolic reorganization. During the initial phase (1 hpt), early SnRK1 activity and an accumulation of specific metabolites, such as succinate, suggest a transient metabolic burst (Figs. 3 and 4). However, by 1 dpt, roots may enter a more energy-conserving state. This latter response is characterized by a pronounced Suc peak (+85.3% respect to 1 hpt) and a simultaneous depletion of glycolytic and TCA cycle intermediates. This pattern is consistent with a coordinated reduction in respiratory flux and ATP demand under stress (Munns & Tester, 2008; Zhang *et al*., 2024). Crucially, at 1 dpt features the uncoupling of the canonical Suc–Tre6P nexus: as Suc peaks, Tre6P declines (−43.4%), and SnRK1 activity drops by 35.6% relative to 1 hpt. Accordingly, PCA analysis confirmed that SnRK1 activity was inversely correlated with the metabolic reprogramming observed at 1 dpt (Fig. 7A). Correlation analyses further capture this unique stress signature, revealing significant negative relationships between Suc and both SnRK1 (r = −0.88, P<0.0002) and Tre6P (r = −0.63, P<0.02). Therefore, our data supports a model in which the canonical coupling between sucrose, Tre6P, and SnRK1 is transiently disrupted during the early phase of salt stress. Rather than reflecting a direct causal breakdown, this uncoupling likely emerges from the integration of multiple stress-derived signals that override the canonical carbon-dependent regulation of SnRK1. Several non-mutually exclusive mechanisms may explain this transient uncoupling. First, alternative sugar phosphates or metabolic intermediates could modulate SnRK1 activity independently of Tre6P (Fichtner et al., 2020; Avidan et al., 2023; Eom et al., 2024). Second, the early oxidative burst, evidenced by MDA, increase under salinity may transiently alter SnRK1 activity through redox-sensitive residues. Third, hormonal signals such as ABA may contribute to the rapid reprogramming of energy signaling pathways. While our data do not allow us to discriminate between these mechanisms, they collectively support the existence of a multi-layered regulatory framework. Ultimately, this metabolic downscaling at 1 dpt effectively decreases structural biosynthesis, evidenced by reduced UDP-Glc and Gal1P levels, thereby prioritizing cellular survival over active growth (Baena-González & Lunn, 2020; Tong et al., 2025).

As stress progresses into the late response phase (1 to 7 dpt), roots exhibit a distinct metabolic profile indicative of acclimation. Tre6P progressively recovers to near-control values, associated with a reactivation of SnRK1 (+40.71% at 7 dpt compared to 1 dpt). During this phase, sustained carbon reallocation away from structural anabolism promotes the synthesis of compatible solutes. The marked accumulation of proline, decrease of soluble proteins, alongside increased FRAP, underscores a restored redox and osmotic balance (Shabala et al., 2016; Li et al., 2023). Evidence has established that the plant SnRK1 catalytic subunit is directly modulated by the cellular redox state through highly conserved cysteine residues (e.g., Cys130 and Cys174 located near to the activation T-loop) (Wurzinger et al., 2018). While the sudden oxidative burst during the early response of salt stress, evidenced by MDA increase at 1-3 hpt, could regulate and temporarily attenuate kinase activity (−31.6 % lower activity from 1 hpt to 1 dpt), the later stabilization of the antioxidant capacity, evidenced by the elevated FRAP at 3 dpt, likely restores the optimal reducing environment required for full SnRK1 functioning. Furthermore, active SnRK1 has been shown to directly phosphorylate respiratory burst oxidase homologues (RBOHs), driving localized ROS signaling that reinforces stress acclimation and nutrient uptake (Zheng et al., 2024). Together, these mechanisms ensure sustained activation of the SnRK1 signaling cascade, coordinating long-term metabolic adaptation under prolonged salinity. Notably, at 7 dpt, both Suc levels and SnRK1 activity increase in parallel, an unexpected outcome given the canonical inhibitory effect of sugars on the kinase. A likely explanation lies in subcellular compartmentation: in the highly vacuolated cells of alfalfa roots, bulk Suc is probably sequestered into the vacuole as an osmoprotectant, masking the discrete dynamics of the smaller nucleo-cytosolic sugar pools that directly govern SnRK1 (Ramon *et al*., 2019; Lando *et al*., 2026). This spatial decoupling might allow the persistent Suc–SnRK1 axis to coordinate osmotic defense while sustaining essential energy signaling for long-term viability (Sarkar & Sadhukhan, 2022; Morales-Herrera et al., 2024).

This biphasic metabolic reprogramming underlies the observed root architectural plasticity, as LR formation is an energy-demanding process tightly controlled by the SnRK1–Tre6P network (Morales-Herrera *et al*., 2024). The role of SnRK1 in LR development depends heavily on context, resolving apparent contradictions in the literature. Under optimal conditions, SnRK1 acts as a repressor of root branching (Morales-Herrera *et al*., 2023). However, under environmental stress, SnRK1 switches to a promoting role. For instance, SnRK1 activation facilitates LR initiation by phosphorylating bZIP63, which subsequently activates the auxin response factor ARF19 (Muralidhara *et al*., 2021), an auxin-mediated pathway conserved in crops like peach (Zhang *et al*., 2020; Wu *et al*., 2025). In our alfalfa model, the early activation of SnRK1 could be coupled with its intricate interplay with ABA-mediated quiescence pathways (Belda-Palazón *et al*., 2020), likely mediating the transient suppression and later recovery of LR growth. Upstream, the salt-induced depletion of the Tre6P carbon-status sensor at 1 dpt likely releases the inhibition on SnRK1. As metabolic homeostasis recovers during the late response, the coordinated action of the Tre6P–SnRK1 network strategically positions new LRs to optimize water and nutrient uptake. Taken together, these patterns define a ‘suppression-recovery’ sequence in roots: an initial strategic downscaling (reduced flux, low Tre6P, transient SnRK1 attenuation, MDA increase) followed by a sustainable, stress-adapted state (SnRK1 reactivation, Tre6P recovery, osmoprotectant accumulation). Interestingly, a similar phase-dependent kinetic has been documented for H□-ATPase activity in Arabidopsis roots (Bose et al., 2014), suggesting a conserved regulatory motif. From an agronomic perspective, the spatial and temporal modulation of the SnRK1–Tre6P–Suc nexus represents a critical trait for maintaining root vitality. The capacity of alfalfa to accumulate sucrose for osmotic defense while simultaneously sustaining SnRK1-driven energy signaling prevents metabolic collapse during the transition to acclimation. Identifying these metabolic checkpoints opens new avenues for precision breeding: enhancing carbon use efficiency and developing forage cultivars that balance high biomass production with survival in saline soils.

## Supplementary data

Fig. S1. Experimental set-up

Fig. S2. Ion concentration in roots of alfalfa

Fig. S3. Effect of salinity on AMPK phosphorylation and SnRK1 marker gene expression in alfalfa root.

Table S1. Primer list

## Acknowledgments

We thank Monica Cornacchione and Ariel Odorizzi for providing alfalfa seeds and Technician Paola Suarez for technical assistance. Prof John Lunn for metabolomic analysis collaboration.

## Author contribution

MR conceptualization. GB and MR designed and performed the experiments. GB, RP methodology. RF performed the LC-MS/MS measurements and analyzed the data. GB, RP and MR formal analysis. GB and MR created and edited the final Figures. GB, RP, MR writing - original draft. GB, RP, RF, MR writing - review & editing. MR: supervision; MR: funding acquisition.

## Data availability

The data that support the findings of this study are available from the corresponding author upon reasonable request.

## Abbreviations

LR: lateral root
PCA: Principal component analysis
PR: primary root
SnRK1: Sucrose non-fermenting kinase 1
Suc: sucrose
TCA: Tricarboxylic acid
Tre6P: Trehalose 6-phosphate
VIP: Variable Importance in Projection

